# Single-cell sequencing of tumor-associated macrophages in a *Drosophila* model

**DOI:** 10.1101/2023.06.17.545411

**Authors:** Dilan Khalili, Mubasher Mohammed, Martin Kunc, Johan Ankarklev, Ulrich Theopold

## Abstract

**Introduction:** Tumor-associated macrophages may act to either limit or promote tumor growth, yet the molecular basis for either path is poorly characterized.

**Methods:** We use a larval *Drosophila* model that expresses a dominant-active version of the Ras-oncogene (Ras^V12^) to study dysplastic growth during early tumor progression. We performed single-cell RNA-sequencing of macrophage-like hemocytes to characterize these cells in tumor- compared to wild type larvae. Hemocytes included manually extracted tumor-associated- as well as circulating cells.

**Results and discussion:** We identified 5 distinct hemocyte clusters. In addition to Ras^V12^ larvae we included a tumor model where the activation of effector caspases was inhibited, mimicking an apoptosis-resistant setting. Circulating hemocytes from both tumor models differ qualitatively from control wild-type cells – they display an enrichment for genes involved in cell division, which was confirmed using proliferation assays. Split analysis of the tumor models further reveals that proliferation is strongest in the caspase-deficient setting. Similarly, depending on the tumor model, hemocytes that attach to tumors activate different sets of immune effectors – antimicrobial peptides dominate the response against the tumor alone, while caspase inhibition induces a shift toward members of proteolytic cascades. Finally, we provide evidence for transcript transfer between hemocytes and possibly other tissues. Taken together, our data support the usefulness of *Drosophila* to study the response against tumors at the organismic level.

## 1 Introduction

When cellular homeostasis is impaired, affected cells may limit the damage by inducing either cellular arrest or cell death. Best characterized in this context is the tumor suppressor p53, which – depending on the amount of DNA damage or other forms of cellular stress, induces cellular senescence, apoptosis, or alternative forms of cell death (1–3). Collectively or individually, these responses will prevent malfunction at the tissue/organ and organismic levels. Notably, this may also prevent the transition from benign to more aggressive forms of tumor growth by eliminating or silencing damaged cells locally in the tumor microenvironment (TME) and at an early stage of tumorigenesis (1). Cell cycle arrest is usually expected to prevent this transition on its own while apoptosis involves the clearance of apoptotic bodies by phagocytic neighboring cells or professional phagocytes such as macrophages, which migrate into the TME (2,3).

While apoptosis thus acts as a tumor suppressor mechanism, tumor cells have been found to evade cell death and in several tumors with poor prognosis, high levels of apoptosis are detected (4). This raises the question of whether apoptosis, both when occurring naturally or as part of anti-cancer treatments is fully beneficial (4).

Tumor progression depends on factors that act both locally in the tumor microenvironment as well as on the communication with tissues in trans, which modulate physiological and immune responses towards tumors. Key actors in the TME are tumor-associated macrophages (TAMs), which may affect tumor progression both positively or negatively. In *vitro* activated macrophages have been roughly classified as either classically activated (M1-type macrophages) or alternative (M2 macrophages, (5)). M1 macrophages are pro-inflammatory and are involved in anti-viral, anti-bacterial, and anti-tumor responses, whereas M2 macrophages are anti-inflammatory, contribute to anti-helminth and tissue repair responses, and are considered pro-tumoral (6,7). Additional *in vitro* and *in vivo* data led to further subdivision, in particular of M2 macrophages, and show that their activation state is only partially reflected by the M1/M2 classification (6,8).

Facilitating the communication within the tumor microenvironment, which includes tumor cells, non-tumor stroma, and TAMs, extracellular vesicles (EVs) have gained increasing attention in recent years (4). EVs include (a) tumor-derived apoptotic fragments, (b) microparticles, which are shed from the cell surface, and (c) exosomes, which are released through the fusion of multivesicular bodies with the cell surface (9). EVs may have tumor and metastasis-promoting capacity both within the TME (4) and systemically. Apoptotic bodies contain nuclear fragments (including DNA and primary transcripts) while the content of microparticles and exosomes is derived from the cytosol and therefore contains mature RNAs. Potential intermediaries of EV-mediated communication include proteins, lipids, and different types of RNA, including long non-coding RNAs (lnc RNAs) (9,10).

Despite lacking adaptive immunity in a mammalian sense, insects possess a highly effective innate immune system. This comprises both humoral and cellular elements. Humoral components are secreted into the hemolymph – the insect equivalent of blood – primarily from the fat body, which fulfills functions of both the mammalian fat body and liver. In *Drosophila melanogaster*, one of the major models for insect immunity, the cellular branch comprises three classes of cells collectively called hemocytes (11): (1) plasmatocytes, which are functionally equivalent to both mammalian macrophages and white blood cells (2) crystal cells which contain prophenoloxidase, the precursor for a key enzyme involved in antimicrobial and wound responses and (3) lamellocytes, which are absent in naive animals but produced in response to wounding and invasion by large intruders such as parasitoid eggs. Recent hemocyte transcriptome profiling at the single-cell level revealed a much higher diversity, in particular among plasmatocytes (11–14).

The interplay between fly tumors and hemocytes was first studied in pioneering work by Pastor Pareja et al (15), where it was shown that upon recognition of damage to the basement membrane, hemocytes are recruited to tumor tissue with tumor-limiting effects. Conversely when in subsequent work tumors were induced in a background expressing a dominant-active version of the Ras oncogene (Ras^V12^) tumor-associated hemocytes (TAHs) were instead involved in a positive feedback loop that involved JNK signaling and promoted tumor growth (16,17). Thus, similar to what is found in mammals, the effects of TAHs appear to be in many cases tissue- and tumor stage-dependent. In previous work, we found that expression of dominant active Ras^V12^ in larval *Drosophila* salivary glands (SGs) induced ductal hypertrophy (18) similar to what is observed in ductal tumors in humans. In SGs, this led to (1) the loss of cellular integrity; (2) nuclear disintegration and caspase activation (3) loss of the SG lumen and of secretory activity (4) damage to the basement membrane (5) induction of fibrotic lesions including activation of the flies’ coagulation system and (6) recruitment of TAHs (18–20). Despite the presence of hallmarks of apoptosis and activation of JNK signaling, SG cells were not eliminated (18,20) although SG cell fragments were released into the hemolymph (18). Surprisingly, forced expression of the antimicrobial peptide Drosomycin (Drs) across whole SGs reverted the majority of tumor-associated phenotypes through negative regulation of JNK signaling (19,20).

Here, to identify genes that are differentially expressed in hemocytes from tumor larvae, we profiled the transcriptome of circulating single-cell hemocytes from wild-type and tumor larvae, as well as from tumor-associated hemocytes (TAHs), which were extracted manually from tumorous salivary glands (Fig 1A). Since we had previously observed strong activation of caspases in Ras^V12^-expressing SGs (20), and in light of the bi-edged nature of caspase activation and apoptosis in cancer (see above), we included larvae where effector caspases were inhibited by the expression of the specific inhibitor p35.

**Figure 1.**
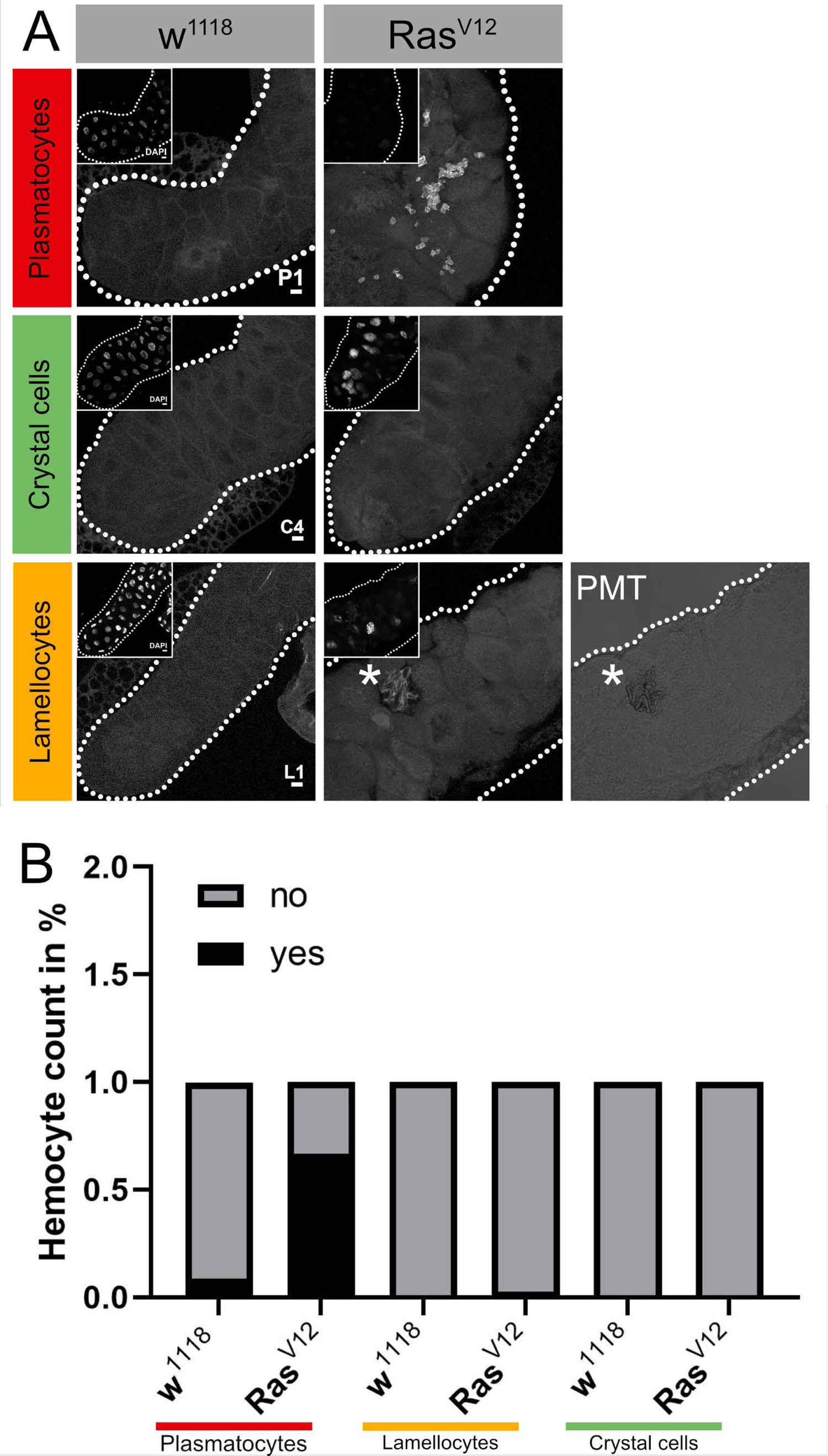
Classification of Tumor-associated hemocytes. **(A)** Tumor-associated hemocytes were labeled using plasmatocyte (P1), crystal cell (C4), and lamellocytes (L1)-specific antibodies. **(B)** The frequency of the three hemocyte classes is shown in the lower part for wild type and Ras^V12^-SGs.

## 2 Material and Methods

### 2.1 Fly strains and sample preparation

*w^1118^, Beadex^MS1096^-Gal4* (Referred here as *Bx*: 8860/Bl), *Beadex^MS1096^-Gal4; EaterDsRed, w^1118^*, *w^1118^;p35* (5072/Bl), *w^1118^;;UAS-Ras^V12^*(4847/Bl), *w^1118^;p35;UAS-Ras^V12^* flies were used in the experiments. Flies were cultured in a 25°C, 12-h dark/light cycle room. Female virgins were collected for 5 days and crossed with respective males on day 7. Progeny larvae were kept as described in (20). Twenty to thirty salivary gland pairs were fixed in 4% PFA for 20 min. Samples for extracellular staining were washed 3 x 10 min with 1x phosphate-buffered saline (PBS).

### 2.2 Immunohistochemistry

Antibodies against P1 (Plasmatocytes, 1:50), L1 (Lamellocytes, 1:50), C4 (Crystal cells, 1:50) were incubated for 1 h at room temperature in PBS and subsequently, washed 3 x 10 min in PBS. The samples were incubated with secondary antibody Anti-Mouse-546 (1:500, ThermoFisher #A11030), DAPI (1:500, Sigma-Aldrich D9542) for 1 hour at room temperature and washed 3 x 10 min with PBS before mounting in Fluoromount-G (ThermoFisher, 00-4958-02).

### 2.3 Proliferation assay

*Drosophila* larva at 120 hours AED was bled into 10 µl of PBS and the sample was incubated for 5 min at room temperature. The sample was fixed in 4% PFA for 20 min and washed 3 x 5 min with PBS. The sample was stained with antibody against pH3 (phosphorylated histone 3, 1:800, Sigma-Aldrich 06-570) and H2 (Hemese, 1:5 kindly provided by Istvan Ando, Szeged) overnight in 4°C. After incubation, the sample was washed 3 x 5 min with PBS and stained with secondary antibodies Anti-Rabbit-568 (1:500, ThermoFisher A-21069), Anti-Mouse-488 (1:250, ThermoFisher A-11001) and DAPI (1:500, Sigma-Aldrich D9542) for 2 h in room temperature. Subsequently, the sample was washed 3 x 5 min with PBS and mounted in Fluoromount-G (ThermoFisher, 00-4958-02). 15 samples per genotype were prepared and photographed using a tile scan at a confocal microscope Zeiss LSM780 (Zeiss, Germany), and the count of cells was evaluated in ImageJ (version 1.53t). Statistics and graph plotting was performed in Prism software (GraphPad Software, 9.3.0, USA).

### 2.4 Image acquisition and analysis

Whole salivary glands were photographed using Zeiss Axioscope II (Zeiss, Germany) microscope and images were exported as .tiff files. The intensity was measured using ImageJ (version 1.53t) and representative pictures were obtained from a confocal Zeiss LSM780 (Zeiss, Germany) microscope. Statistics were performed in Prism software (GraphPad Software, 9.3.0, USA).

### 2.5 Micro-manipulation of attached hemocytes

Ten pairs of salivary glands per genotype were dissected into individual droplets of 200 µl PBS. A fluorescent microscope Leica DMi8 (Leica, Germany) was used to locate the attached hemocytes at the surface of the salivary gland. The hemocytes were collected using micromanipulator TransferMan NK2 (Eppendorf, Germany) with ø 7 µm collection capillary and separated by piezo-vibrator PiezoXpert (Eppendorf, Germany). Single-cell state hemocytes were individually transferred to 2.3 µl of lysis buffer (38) and each sample library was prepared as described below.

### 2.6 cDNA synthesis and library preparation for attached hemocytes

cDNA libraries of single attached hemocytes were generated using a modified version of the Smart-seq2 protocol (38). In short, cDNA synthesis was performed using universal primers, and PCR amplification was carried out over 24 cycles. cDNA products were subsequently purified using CA beads (Sigma; catalog no. 81260) for size selection using 8.8% polyethylene glycol 6000 (PEG 6000) to exclude primer-dimers and nonspecific amplicons with sizes less than 150 bp. Combinatorial indexing via tagmentation was carried out in 96-well plates using 200 pg (measured in a Qubit fluorometer) of amplified cDNA, for a final volume of 10 mL/well. cDNA fragmentation using Tn5 transposase was carried out for 20 min on ice using the Illumina Nextra XT DNA sample preparation kit. Ligation and amplification of adaptors were carried out over 15 cycles in a final volume of 25 mL/well. Primer indices were used in the reaction from Illumina (Nextera index primers i7 and i5, catalog no. FC-131-1001). Tagmented and barcoded amplicons were then purified using CA beads for size selection. Quality control and fragment size distribution of the cDNA libraries were performed on a Bioanalyzer with the Agilent high-sensitivity DNA chip (catalog no. 5067-4626). Concentrations of each sample of cDNA libraries were measured on a PicoGreen 96-well plate NucleoScan fluorometer using a high-sensitivity double-stranded DNA (dsDNA) (HS assay kit; catalog no. Q32851). To perform library dilutions, the average fragment sizes of all cDNA libraries were measured for a final concentration of 2 nM in each sample. Finally, cDNA libraries were pooled and sequenced using Illumina NextSeq with 75-bp paired-end reads.

### 2.7 Cell sorting and cDNA library preparation for circulating hemocytes

Ten larvae per genotype were bled into 500 µl of PBS and the cells were sorted. The sorting was performed with a MoFlo Astrios EQ (Beckman Coulter, USA) cell sorter using a 488 and 532 nm laser for excitation, 100 µm nozzle, sheath pressure of 25 psi, and 0.1 µm sterile-filtered 1 x PBS as sheath fluid. Flow sorting data was interpreted and displayed using the associated software, Summit v 6.3.1.

To test the precision of the adjustments made to center the drop in each well, a colorimetric test mimicking the sort was done based on (39). A 1.5 mg/µl solution of HRP (cat no 31490, ThermoFisher Scientific) with 1 drop of flow check beads (Beckman Coulter, USA) was sorted into each well of an Eppendorf 384-well plate (Cat no 34028, ThermoFisher Scientific). A color change after sorting indicated that the drop hit the sort buffer and that the precision was adequate.

Single hemocytes were sorted directly into a 384-well plate containing 2.3 µl of lysis buffer (Eppendorf twin.tecTM PCR plates) using a CyClone^TM^ robotic arm and at highly stringent single cell sort settings (single mode, 0.5 drop envelope) and cDNA libraries were generated by the Eukaryotic Single Cell Genomics Facility at SciLifeLab, Stockholm using a slightly modified version of Smart-seq2 as previously described (38), but where we used 20 cycles for cDNA amplification. The plate and sample holder were always kept at 4 °C during the sort. After the sort, the plates were immediately spun down and put on dry ice.

### 2.8 Single-cell RNA sequencing

Single-cell libraries were sequenced at the National Genomics Infrastructure, SciLifeLab Stockholm, using the HiSeq2500 platform (Illumina) for 56 bps single-end sequencing. We sequenced a total of 463 (negative controls n=2, per plate).

### 2.9 Mapping, annotation and filtering of low-quality cells

A reference genome *Drosophila melanogaster* (Dm6 v r6.37) was indexed and raw fastq files were used in mapping to the genome using STAR v 2.7.2 (40), and gene expression was measured using featureCounts v 2.0.0 (41), using default settings. expression matrix was filtered to obtain high-quality cells using the following criteria: cells with >5% mitochondrial transcripts (stressed/dead/dying cells), <200 genes (low-quality cells), those expressing more than 4000 features (genes) (potential doublets or triplets) were removed in each replicate and the remaining cells were subjected to subsequent computational analysis.

### 2.10 Normalization, dimensionality reduction and clustering

The main computational analysis of read-count matrices was performed using the Seurat package (v 4.0.3) (42) in R (v 4.1.0). The complete R workflow can be assessed and reproduced in the R markdown (see code availability section). we used the default processing pipeline https://satijalab.org/seurat/v3.2/pbmc3k_tutorial.html. First, count matrices and metadata were loaded. A mitochondrial gene count above 10 % was filtered. Quality filtering was performed and cells with a minimum of 200 genes expressed were kept for further processing. Subsequently, reads were normalized for sequencing depth using the “NormalizeData” function in the Seurat toolkit, selecting the top 2000 variable genes. Thereafter, dimensionality reduction was performed using PCA computing the first 50 PCs. The first 10 PCs from the analysis were then subjected to shared-nearest-neighbor (SNN) inspired graph-based clustering via the “FindNeighbors” and “FindClusters” functions. For modularity optimization, the Louvain algorithm was used, and clustering was performed at a resolution of 0.8 for clustering granularity, resulting in 9 clusters. After clustering, a UMAP dimensionality reduction was performed using the first 10 dimensions of the PCA.

### 2.11 Differential gene expression analysis (DGEA)

Differential gene expression analysis (DGEA) of genes in identified clusters was performed using the function “FindAllMarkers” from the Seurat package (v. 4.0.3). Following the default option of the method, differentially expressed genes for each cluster were identified using a non-parametric Wilcoxon rank sum test. Differentially expressed genes in a cluster were defined by setting initial thresholds above a logarithmic fold-change of 0.5 and being present in at least 25% of the cells belonging to the same cluster. Representative marker genes with an adjusted *p*-value below 0.05 for each cluster were further selected. *p*-values were adjusted using a Bonferroni correction including all genes in the dataset. To find representative marker genes with elevated expression in comparison to the remaining clusters, only positive log fold-changes were considered. For individual analyses such as gene enrichment analysis (see “Gene set enrichment analysis (gsea)”), threshold values for differential gene expressions were modified and will be described in detail in the respective sections of the materials and methods and results. To identify DEGs between specific clusters of interest, the “FindMarkers” function in Seurat was used and the identities were set to the respective clusters of interest. The same thresholds as stated above were used to define DEGs.

### 2.12 Biological pathways and GSEA

To track tests for top functional class enrichment among the global clusters, we used “ClusterProfiler” package v 3.16 (43) tool to conclude the enriched ontology terms as previously mentioned specifying the database “Org.Dm.db” to calculate the top 5 biological pathway enrichment. The gene set enrichment analysis (GSEA) was performed on top differentially expressed genes over the identified clusters in regards to gene expression profiles “Log2FC” as input in cluster profiler v 3.18.1 and ggupset package v 0.3.1 with a *P value* cutoff of 0.05, minGSSsize of 3, maxGSSize of 800, and scoreType of “pos” to estimate for biological process ontology across clusters.

### 2.13 Computational summary

The read alignment and gene count matrix generation were performed as previously described (Methods). The single-cell gene count matrix cells with fewer than 250 UMIs, more than 10,000 UMIs, reads mapping to more than 7000 genes, or more than 10% of read counts mapping to ribosomal genes were excluded. Each single cell transcriptome was mapped to its original time window from which it was extracted by using the RT barcode. We performed standard processing of the data split by experiments as recommended by Seurat v4 documentation including NormalizeData, FindVariableFeatures (with the method set to ‘vst’), ScaleData, RunPCA, RunUMAP (with dims set to 1:50 and n.components to 2), FindNeighbors (with reduction set to UMAP and dims set to 1:2), FindClusters. This version uses the first 10 principal components, along with pN of 0.2 and pK of 0.005. Dimensionality reduction, clustering, and identifying cluster-specific marker genes. The standard Seurat processing pipeline, as described in the previous section, was performed on each non-overlapping inferred age window separately. We found that the default Seurat clustering resolution parameter did not capture the dynamics of the presence of different cell types. For each batch, we clustered the data with a variety of resolutions (from 0.1 to 1.5 in increments of 0.3) and then computed the within-cluster sum of squares (WSS). Visualizing WSS across increasing resolutions we chose the resolution in which there was a visible plateau in the decrease of WSS. Finally, we integrated and clustered the data and from these clusters, we used Seurat’s FindMarkers to iteratively loop through all clusters and identify marker genes.

### 2.14 RNA velocity and lineage interference

To interpret the global transcriptional progression of hemocytes and their cell fate decision, we established the cell continuum of cell differentiation, data layers of unspliced and spliced mRNA for the entire data generated with Velocyto CLI (v.0.17.17) according to the CLI usage guide (Velocyto run-smartseq2). The output loom files were combined using “loompy”. The merged loom file was imported into the scVelo package (v1.0.6) (44,45). The unspliced and spliced mRNA counts of cells from clusters C0-C4 were extracted. We used the “merged.utilis” function in the scVelo pipeline, where cells with low pre-mRNA counts were removed as part of the filtering. In short, the gene-specific velocities are obtained by fitting a ratio between unspliced and spliced mRNA abundances and then computing how the observed abundances change from those observed in a steady state. The ratio of ‘spliced’, ‘unspliced’, and ‘ambiguous’ transcripts were calculated and data were pre-processed using functions for detection of minimum count number, filtering and normalization using “scv.pp.filter_and_normalise” and followed by “scv.pp.moments” function. The gene-specific velocities were then calculated using “scv.tl.velocity” with mode set to “deterministic” and “scv.tl.velocity_graph” function to generate velocity graph, and visualization using “scv.pl.velocity_graph” function. In addition, we used the “scv.tl.recover_latent_time” function to infer a shared latent time from splicing dynamics and plotted the genes along the time axis.

### 2.15 Data availability

The raw processed data generated for this study have been deposited on Zenodo repository and can be accessed via [https://zenodo.org/deposit/7997643] and interactively on [https://mubasher-mohammed.shinyapps.io/Sc-drosophila/]. The custom code scripts used to analyze data for this study are available at [https://github.com/ANKARKLEVLAB/Single-cell].

## 3 Results

### 3.1 Single-cell profiling of tumor-associated hemocytes reveals transcriptional heterogeneity

We initially characterized the population of tumor-associated hemocytes (TAHs) that attach to Ras^V12^ salivary glands (SGs) and found they consist mainly of plasmatocytes and the occasional lamellocyte (**Figure 1A-B**). To identify genes that are differentially regulated in hemocytes (DEGs) from tumor larvae we profiled the transcriptome of single circulating hemocytes from wild-type and tumor larvae as well as from tumor-associated hemocytes (TAHs), which were extracted manually from tumorous SGs. Additionally, to characterize the contribution of effector caspases to tumor progression we analyzed larvae in which the effector caspase inhibitor p35 was co-expressed with Ras^V12^: p35;Ras. We had previously shown that – similar to Drs - expression of p35 in Ras-SGs restores the spherical nuclear shape that is disturbed upon sole Ras-expression although SGs from both combinations show the same size increase as Ras^V12^ larvae (20) and (**Figure 2A-D, quantified in E**). In contrast to Drs (20), co-expression of p35 still led to hemocyte recruitment (**Figure 2A’-D’ quantified in F**). This allowed us to compare tumor-associated hemocytes (TAHs) both between wild-type and tumor larvae as well as in settings with and without active caspases downstream of JNK-signaling. TAHs as well as circulating hemocytes were selected for expression of the plasmatocyte-specific marker Eater (**Figure 3A**, see Material & Methods).

**Figure 2.**
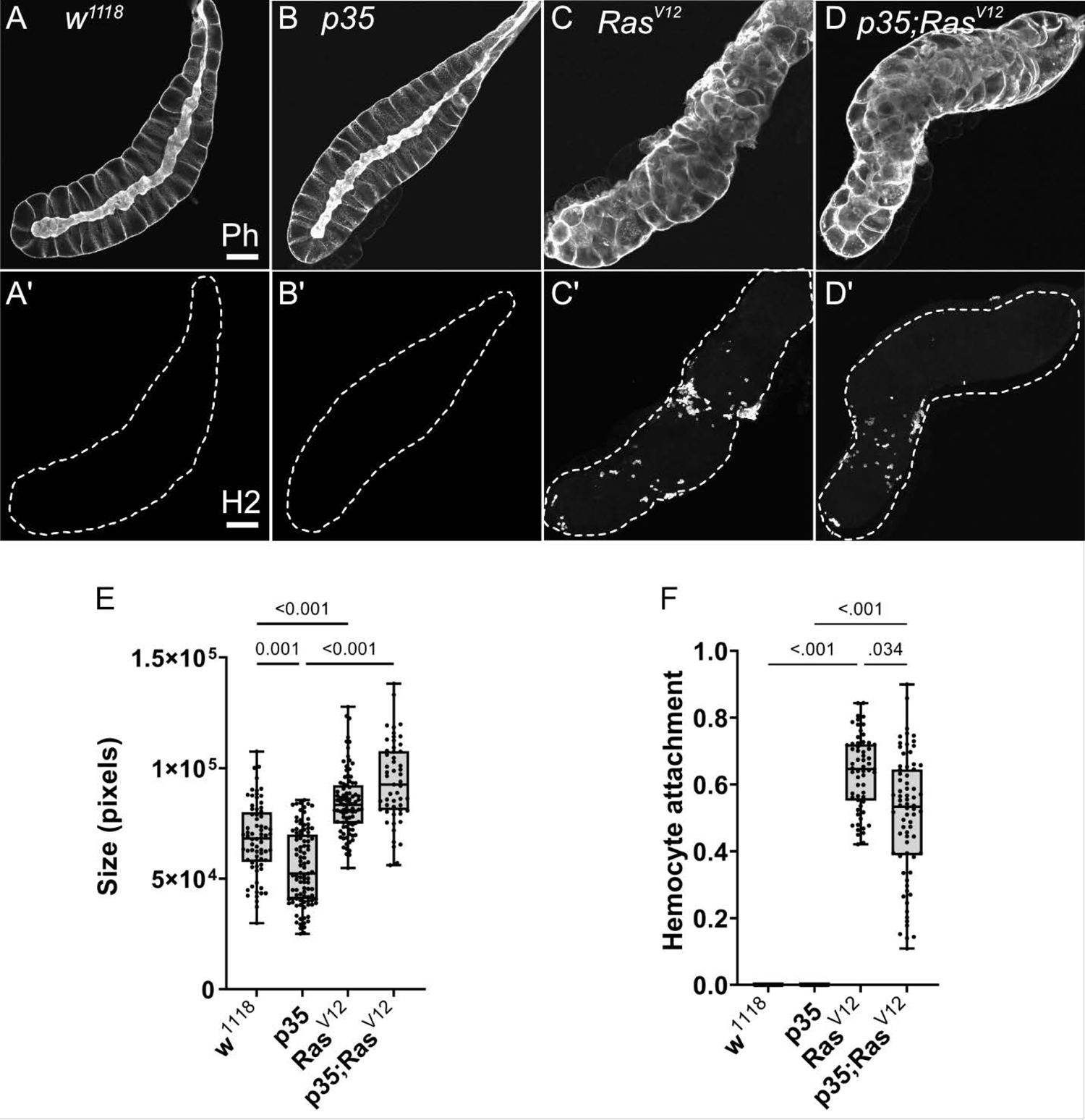
Characterization of SGs from the used genotypes. **(A-D)** SGs from wild type (w^1118^), p35 expressing larvae, Ras^V12^ larvae and p35;Ras^V12^ larvae were stained with phalloidin and a plasmatocyte-specific antibody (Hemese, H2, **A’-D’**). SG size for the different genotypes was quantified shown in **E** and hemocyte attachment in **F**. Whisker length min to max, bar represent median. P-value quantified with ANOVA.

**Figure 3.**
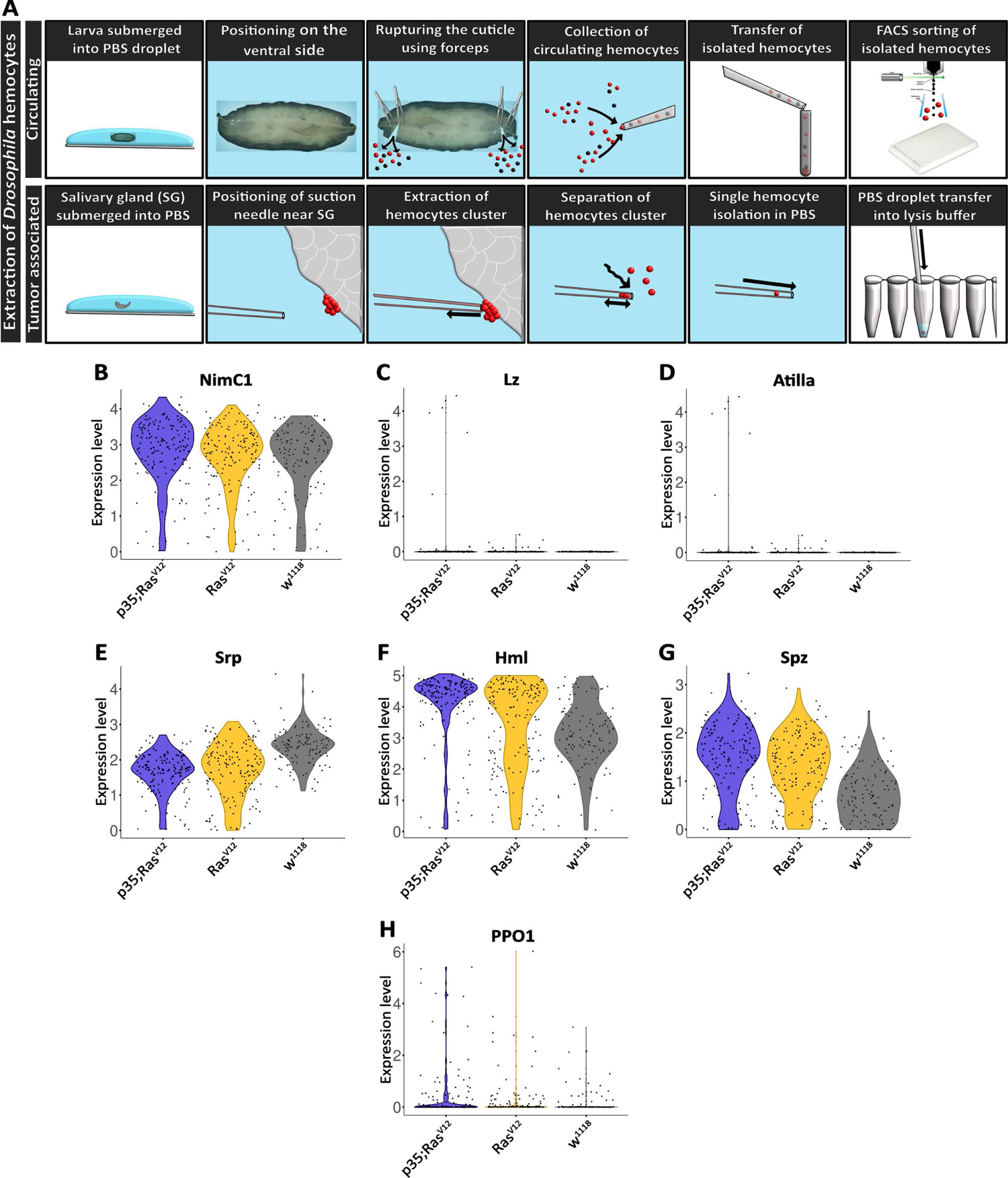
Extraction and initial characterization of circulating hemocytes and TAHs from wild type and tumor larvae. **(A)** Upper panel: circulating hemocytes were extracted from larvae and individualized using cells sorting. Lower panel: TAHs were extracted from salivary glands using capillary suction, individualized and processed for single-cell RNA sequencing. See text for further details. **(B-H)** Classification of single hemocytes confirms their plasmatocyte identity and the absence of lamellocytes (Atilla) or crystal cell (Lozenge) markers.

Collectively, hemocyte transcriptomes displayed additional plasmatocyte markers indicating successful purification of these macrophage-related cells (**Figure 3B-H**). Plasmatocyte markers include Nimrod C1 (NimC1), serpent (srp), Hemolectin (Hml), and spätzle (spz), several of which have been used as pan-plasmatocyte markers and more recently as markers for specific plasmatocyte subpopulations (21). As expected, the crystal cells and lamellocytes markers lozenge (lz) and Atilla, respectively; were not detected (**Figure 3C-D**). Crystal cell prophenoloxidase 1 (PPO1) was detected in a few cells (**Figure 3H**) see also below.

Clustering analysis revealed true variation between the single cell transcriptomes for the identified 5 clusters (at p<0.05 and a difference in expression >2 if not stated otherwise; (**Figure 4A-C, middle panel**), three of which included tumor cells (clusters C0, C2 and C3 (**Figure 4C-D**). Cluster C2 overlaps largely with TAHs (**Figure 4A**). Gene Ontology analysis for biological processes of up-regulated genes in the clusters highlighted several gene sets that overlapped between clusters but also some cluster-specific sets (**Figure 4E**). The latter category included genes involved in cell motility (cluster C1), translation (C0) protein folding (C4) and immune response genes for cluster 2 (attached cells) and several categories specific for cell division (C3, circulating cells from both tumor models). Taken together, despite the technically limited number of cells compared to other studies (21), we identified subpopulations of plasmatocytes whose signatures differed significantly depending on genotype (tumor versus wild type) and cell status (attached versus circulating).

**Figure 4.**
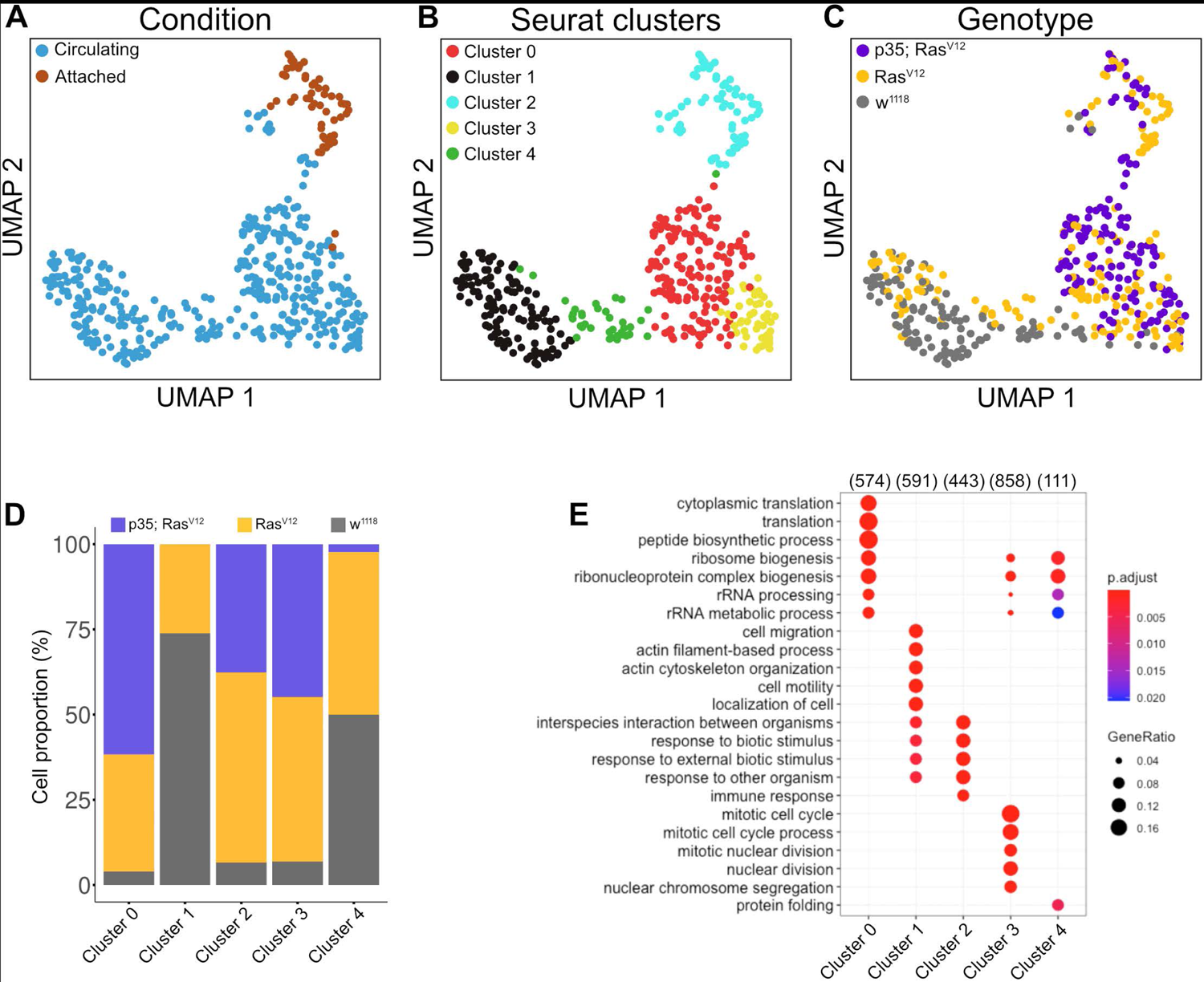
Clustering analysis of hemocytes. Circulating single cells (Genotype: w^1118^, Ras^V12^ and p35;Ras^V12^) were processed using Smart-seq2. For manually extracted TAHs (Genotype: Ras^V12^, p35;Ras^V12^) libraries were generated according to the Smart-seq2 protocol. **(A)** UMAP projection of single-cell transcriptome integrated datasets overlaid with batch labels. **(B)** Louvain clustering analysis based on the first 10 PCs shows the biological variation of the data and cell communities’ assignments. **(C)** UMAP projection overlaid with single-cell specific genotypes. The global transcriptome similarities and differences were assessed based on k-nearest neighbors (kNN) force-directed graph with a true signal variation of the single-cell transcriptome on the integrated datasets. **(D)** Relative frequency of w^1118^, Ras^V12^, and p35Ras^V12^ hemocytes within the five identified clusters**. (E)** Gene set enrichment analysis (GSEA) (Biological processes) of up-regulated genes in a minimum of 25 % of the cells in each cell community with a cut-off 0.25 log fold change threshold, comparing different cell communities (clusters) and highlighting the various biological processes’ heterogeneity. Color scale indicating the corrected p-values where blue is less significant, and red is highly significant. Black circular dots indicate the gene ratio in comparison to the universal background gene list. Numbers in brackets indicate the gene numbers overlapping the ontology terms for a specific cluster.

### 3.2 The expression of signature genes for the different clusters differs between tumor models

To obtain more detailed insight into the gene categories that specified the clusters, we identified the genes whose expression significantly specified the clusters (**Tables S1-S5**). In parallel, we compared expression strengths between genotypes focusing on the three populations that comprised cells from the tumor models (clusters 0, 2, and 3 in **Figure 4D and Figure 5A-C**). For attached hemocytes (C2), the list of the most strongly expressed genes varied between the two tumor models (**Figure 5A**). To follow this up we scanned cluster 2-specific genes for gene-set enrichment and identified two enriched categories: (1) pupal adhesion and (2) immune response (**Figure S1A-B**). Pupal adhesion was due to the presence of genes that code for salivary gland secretions (**Figure S1B**). Since these genes are hardly expressed outside salivary glands (FlyAtlas: (22)), we interpret their presence in hemocytes as being passenger transcripts (23), i.e., transcripts that have been taken up by TAHs by way of cellular fragments released from the glands. Alternatively, SG transcripts may have been co-purified together with TAHs during extraction. This latter explanation implies that expression of p35 partially prevented the release of SG fragments from Ras^V12^ SGs in line with its anti-apoptotic function. The second category (immune genes) contains genes that are most strongly (although not mutually exclusively) expressed in either tumor model (**Figure S1A**). Taken together, we show that hemocytes in both tumor models differ qualitatively from hemocytes in control larvae and that inhibition of effector caspases in tumors reveals additional quantitative differences.

**Figure 5.**
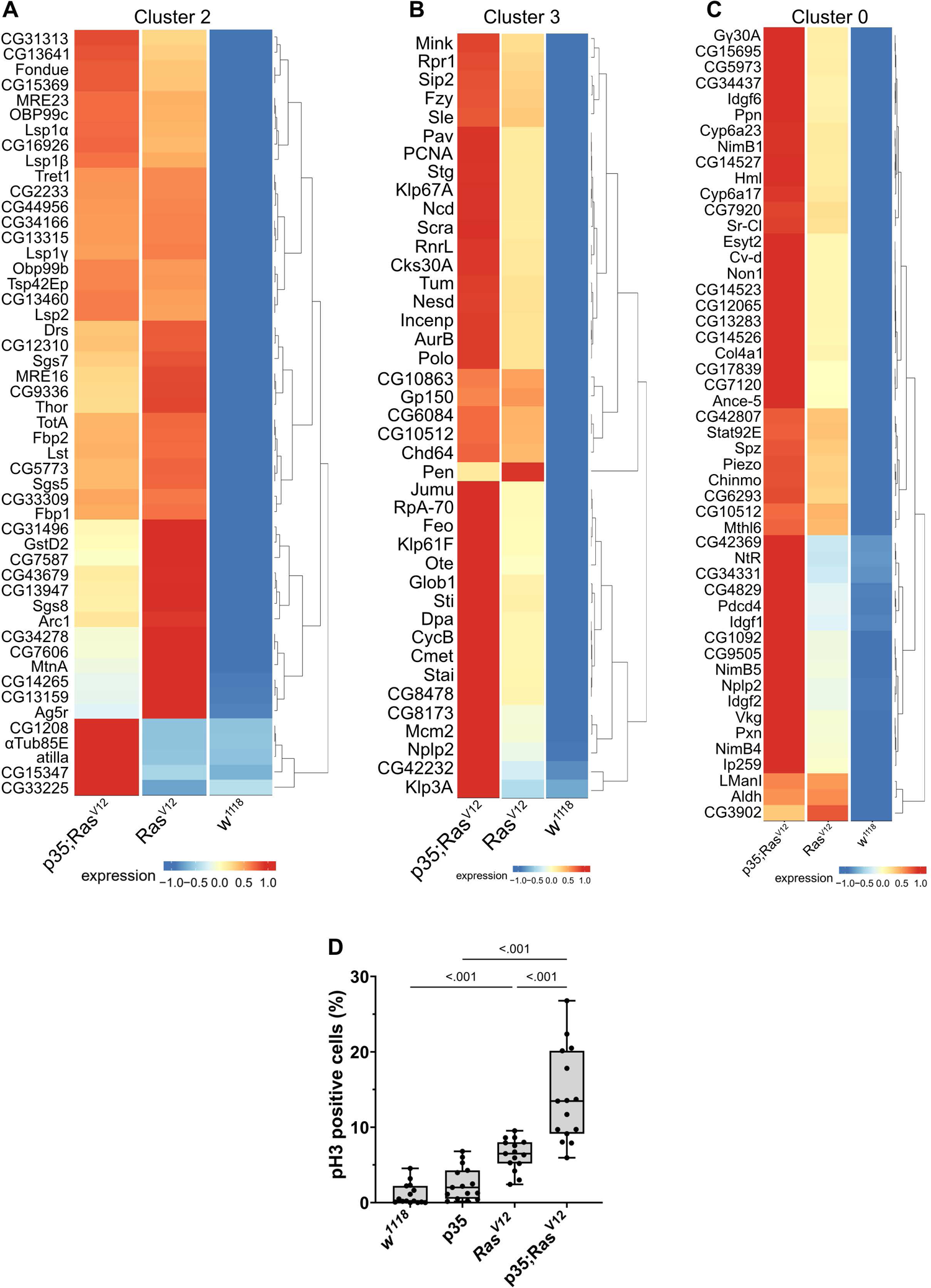
Contribution of the three genotypes to clusters 2 (A), 3 (B) and 0 (C) Expression intensity was analyzed for the most significantly enriched genes in the respective clusters using the visualization tool created for this project (see: https://mubasher-mohammed.shinyapps.io/Sc-drosophila/). **(D)** The enrichment of genes involved in cell division in cluster 3 (see also Figure 4E) was functionally confirmed using anti-phospho Histone 3 labeling to detect proliferating cells. Whisker length min to max, bar represent median. P-value quantified with ANOVA.

### 3.3 The response towards tumors differs between the two models

Generally, TAHs in the Ras model show strong expression of AMPs including Dpt(Diptericin)A and B, Att(Attacin)C, and Cec(Cecropin)C. In contrast p35;Ras^V12^ TAHs show higher levels of all three phenoloxidases PPO1-3 (**Figure S1A**). This is in line with the previously identified function of the TGF-like protein dawdle (*daw*), which promotes AMP activation (24) and indeed, *daw* clusters with AMP expression in our hands (**Figure S1A**). A second TGF member (*dpp*) with immune-regulatory function (24) is less expressed in tumors compared to wild-type hemocytes. Similarly, the M1 marker iNOS is more strongly expressed in p35;Ras^V12^ TAHs while the M2 marker Arginase (Arg) peaks in Ras TAHs.

For both clusters 0 and 3 (**Figure 5B-C**), the same set of genes was more strongly induced in circulating tumor compared with circulating wild-type hemocytes although the majority showed stronger expression when effector caspases were repressed (p35;Ras^V12^) compared to the tumor-only (Ras) model. Gene set enrichment analysis fully confirmed the enrichment for genes involved in cell division (**Figure 4E**) in cluster 3 (circulating cells from both tumor models) but failed to deliver significant returns for the category “Biological process” for cluster 0. In line, the quantification of hemocyte proliferation fully confirmed an increase in cell divisions in both tumor models including the additional increase in the p35;Ras^V12^ model (**Figure 5D**). This may contribute more to the differentiation of TAHs rather than circulating cells, which show similar counts (Hauling et al., 2014).

Finally, when searching for pathways compatible with expression in the five clusters using the Reactome database (https://reactome.org), an enrichment for genes involved in neutrophil degranulation was found for cluster 0 (**Figure S1C**), which when using a split analysis also indicated that this pathway was more strongly expressed in p35;Ras^V12^ hemocytes. Pseudotime analysis using RNA Velocity (**Figure 6A-D**) indicated that TAHs and circulating hemocytes from both tumor models follow distinct pathways although due to the restricted number of TAHs, more rare intermediate population may have been missed.

**Figure 6.**
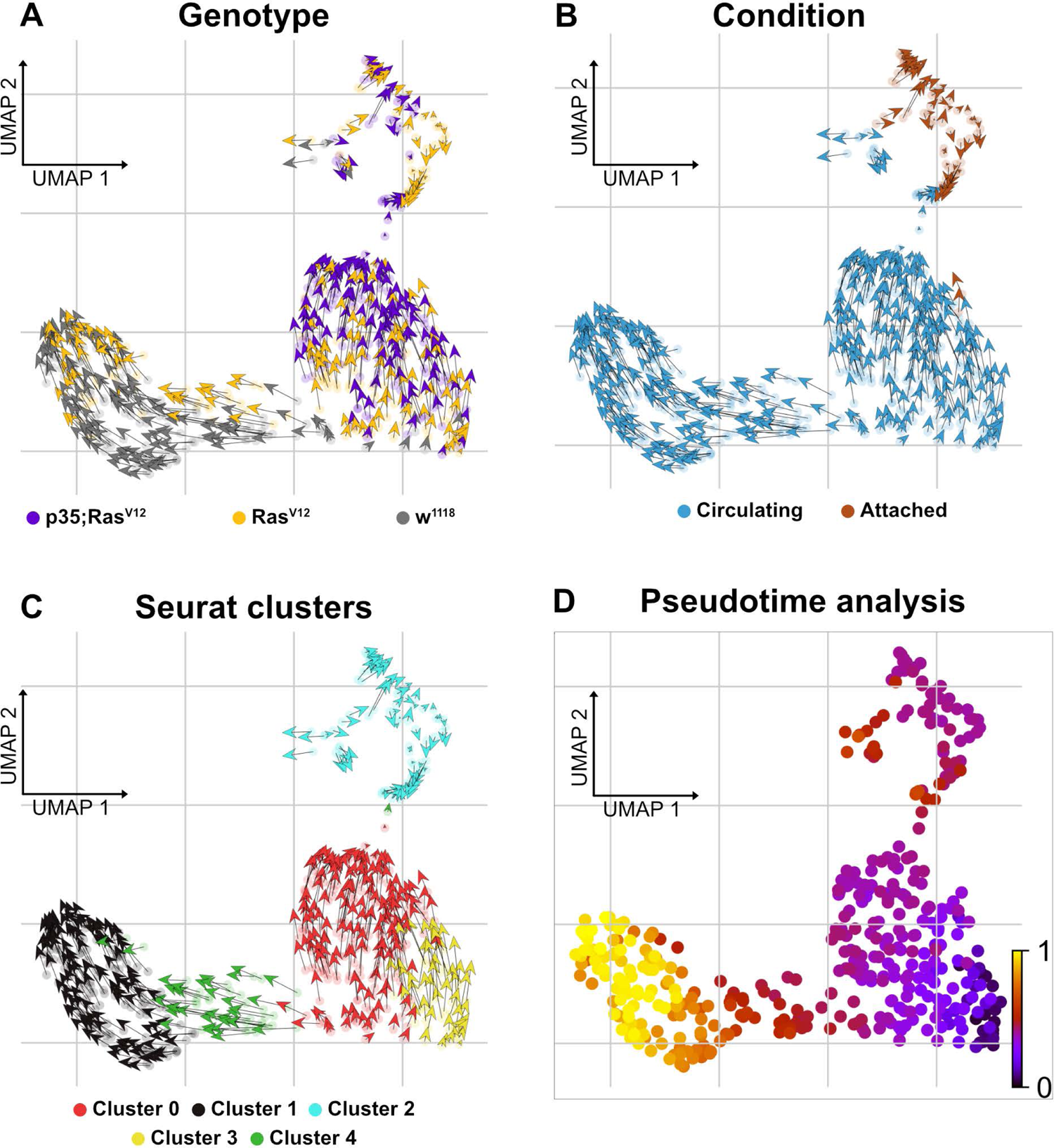
Delineating splicing kinetics through generalizing RNA velocity to cell population clusters. **(A)** Projecting velocities across the wild type, Ras^V12^, and p35;Ras^V12^ hemocytes. (**B)** Across conditions (attached and circulating)**. (C)** Identified Seurat clusters**. (D)** Pseudotime inference of integrated datasets shows the root cells composed mainly of cluster 3 cells (mixture of p35;Ras^V12^ and Ras^V12^ larvae) and trajectory depicting pseudotime units assignment with terminal state composed mainly of cluster 1 (wild type and Ras^V12^ larvae).

### 3.4 Transcript transfer between hemocytes and other tissues

Recently, we and others have characterized the activation of crystal cells and the subsequent release of leaderless PPO2 into the extracellular environment via cell rupture (25,26). Functionally, crystal cell activation bears similarities to pyroptotic cell death in mammals, including its dependence on caspase activity, which can be inhibited by p35 (27). While our findings explained how PPO2 was released into the hemolymph, the mechanism for secretion of PPO1, which also lacks a signal peptide remained obscure. Since we found that PPO2 was enriched in TAHs (**Figure 7A, S1A**), we wondered which secretion mechanism led to their spread to other plasmatocytes. In line with crystal cell rupture and similar to passenger transcripts from SGs, the mRNA encoding PPO2 was detected in plasmatocytes both as a mature (spliced) form and as a non-spliced (nuclear) immature mRNA (**Figure 7C, D**). This indicates that phagocytic hemocytes had access to both the nuclear and the cytosolic fraction of ruptured crystal cells. In contrast, PPO1 transcripts were only detected in their mature form (**Figure 7B**). We conclude that PPO2 transcripts in TAHs originate either from pyroptotic crystal cells or TAHs which have acquired crystal cell characteristics, while PPO1 mRNA originates from live crystal cells.

**Figure 7.**
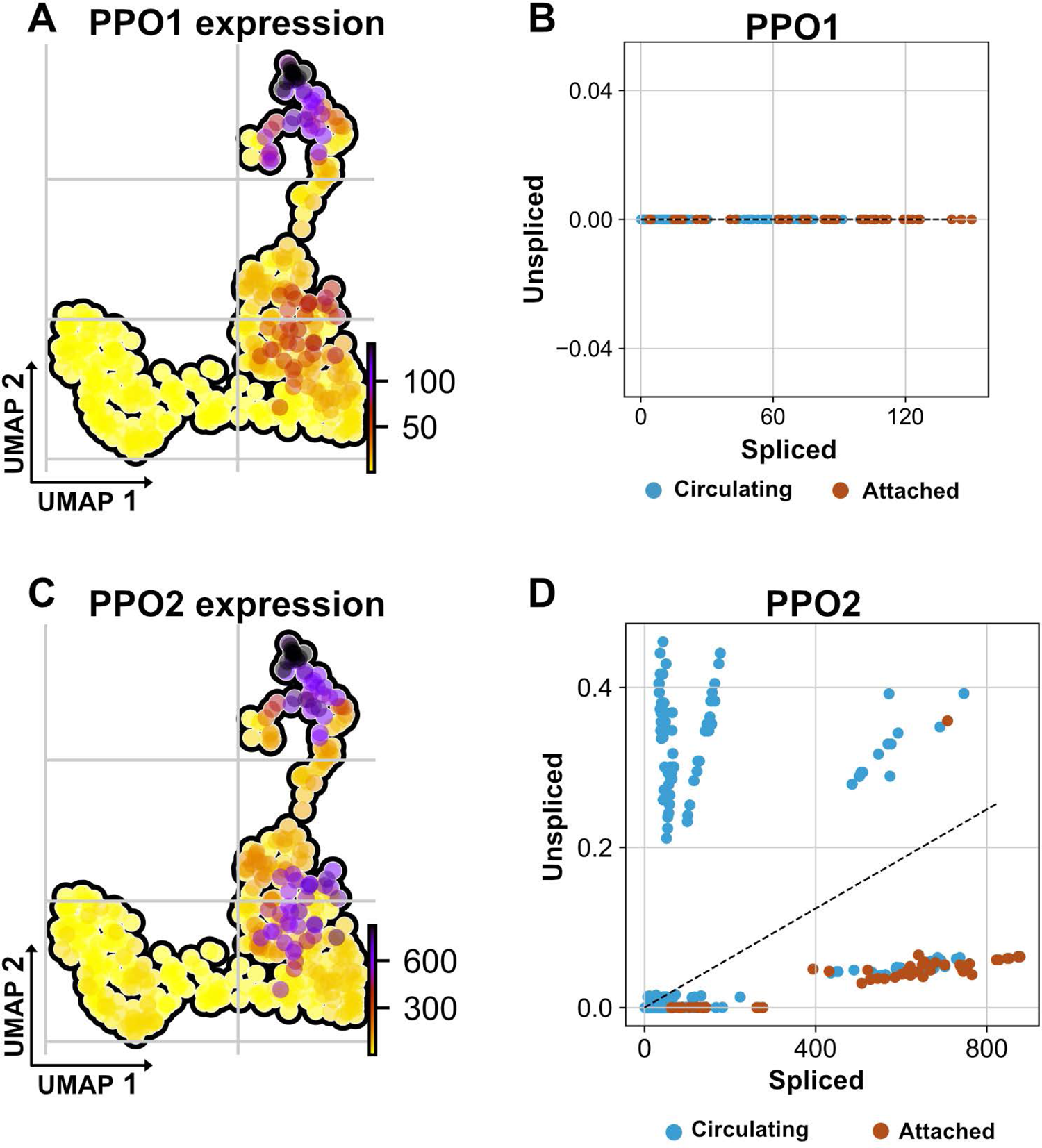
Non-autonomous distribution of crystal cell derived prophenoloxidase transcripts. **(A)** UMAP visualizes transcripts analysis showing the strongest presence of mature (spliced) mRNA of crystal cell-derived prophenoloxidase 1 (PPO1) across attached hemocyte TAHs condition corresponding to Ras^V12^ larvae. **(B)** scatter plot shows the absence of immature transcripts (unspliced) reads across conditions (attached and circulating). **(C)** UMAP visualizes transcripts depicting cell-derived prophenoloxidase 2 (PPO2) in attached hemocyte TAHs condition. **(D)** scatter plot indicates the exclusive presence of mature and immature (spliced vs unspliced) PPO2 transcripts in plasmatocytes (higher ratio of unspliced across circulating condition), in contrast to PPO2 which is released from crystal cells through cell rupture and subsequently taken up by plasmatocytes most likely through phagocytosis.

TAHs also contained transcripts that – according to FlyAtlas – are strongly expressed in fat bodies. When checked for the presence of spliced and unspliced versions we found most of them displayed both, indicating genuine expression in hemocytes, which was strengthened by the data from Tattikota et al who detected them in hemocytes too (21).

In addition to PPO1, 3 transcripts (UDP-galactosyltransferase activity, CG 13962, and the transcription factor Relish) were detected only in their mature form indicating a possible origin outside TAHs.

## 4 Discussion

To our best knowledge, we provide the first transcript profile of tumor-associated hemocytes (TAHs) – the invertebrate equivalent of mammalian tumor-associated macrophages (TAMs). Relying on a molecularly induced early stage of tumor progression that affects the salivary glands (18,20), we find that TAHs display some features of mammalian M2-like macrophages, which have been implicated in regenerative processes. These include the presence of members of the chitinase-like proteins (IDGFs in *Drosophila*), which are amongst the most abundant proteins in activated macrophages and are used as markers for M2 macrophages (28) (**Figure S1C**). Notably, TAHs show a clear signature of immune activation, which includes several antimicrobial peptides as well as members of clotting systems including IDGF3, Fondue, and phenoloxidases (**Figure S1A**). Although both effector branches are activated in the TAHs from both tumor models, there are differences (**Figure S1A**): AMPs appear to dominate the TAH response in Ras-SGs whereas clotting factors are more strongly expressed in p35;Ras^V12^ SGs. AMP induction may be in line with the presence of SG (passenger) transcripts in the Ras-alone model **(Figure 8, right part)** and may serve to degrade tumor fragments during efferocytosis including cytosolic and nuclear parts. In contrast, inhibition of caspases appears to activate immune reactions that are more akin to the formation of mammalian granulomas (**Figure 8, left part).** Bifurcation in hemocyte differentiation is reminiscent of the division of labor between *Drosophila* hemocytes shown previously which depend on two members of the TGF family (the BMP-like member Dpp and the Activin-like member Dawdle, (24)). In line with Dpp’s role in suppressing antimicrobial responses, we find that two receptors for BMP-like TGFs (thickveins; tkv and saxophone; sax) are strongly expressed in p35:Ras^V12^ hemocytes but not in Ras or control hemocytes (**Figure S2**). While the difference in responses between Ras larvae and larvae that co-express the caspase inhibitor p35 may be explained by a lack of apoptosis downstream of caspases, we prefer alternative explanations that include non-apoptotic form of cell death or non-apoptotic functions of caspases (29).

**Figure 8.**
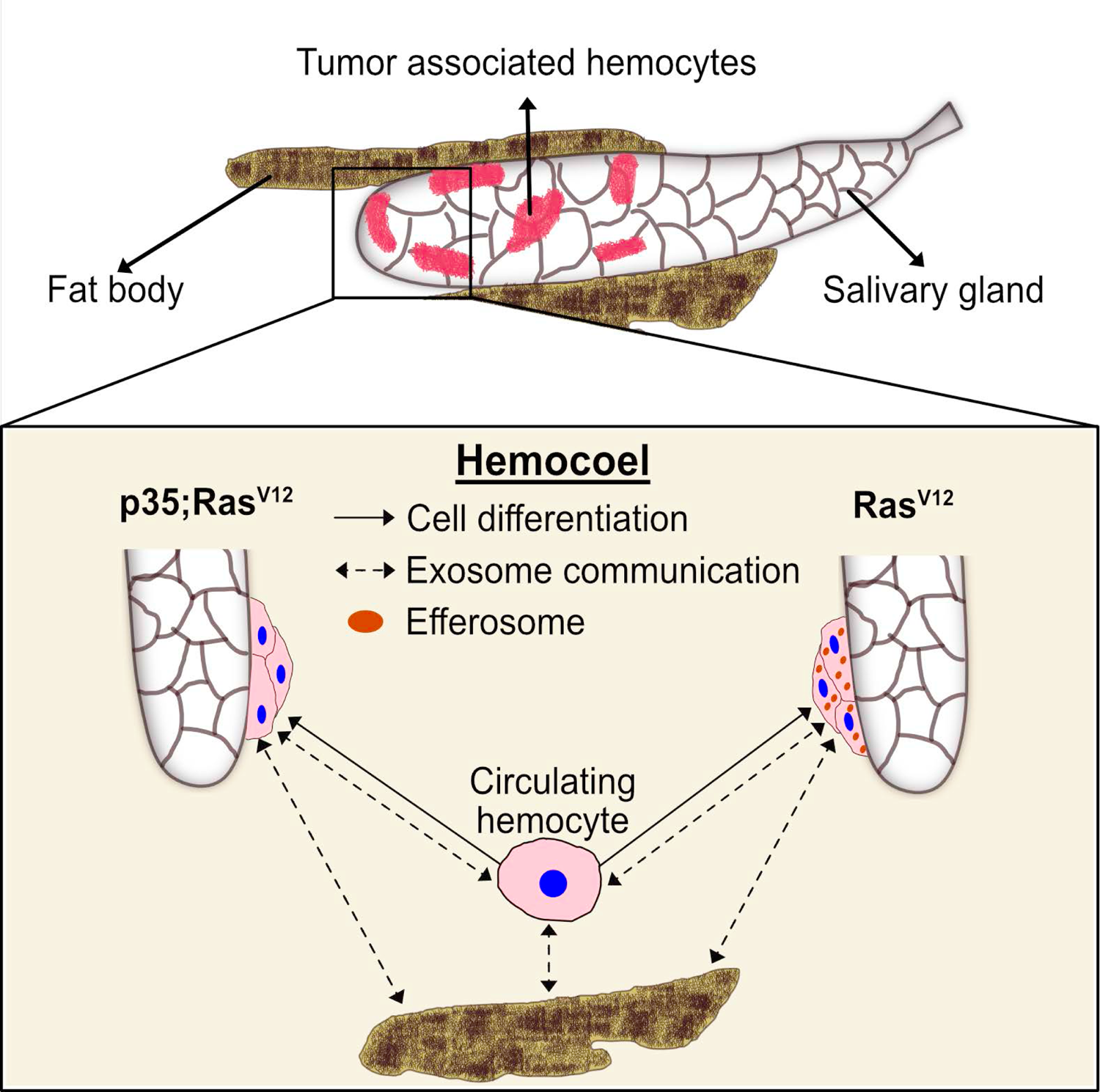
A proposed model for the differentiation and distribution of hemocyte transcriptomes. Hemocyte reactions towards SG dysplasia are schematically depicted for the Ras^V12^ model (right part) and after inhibition of effector caspases by p35 (left part). Communication via exosomes (dashed arrows) and transdifferentiation (solid arrows) are indicated. Efferosomes are present in Ras^V12^ model as result of efferocytosis of apoptotic bodies from hyperplastic salivary gland. See text for further details.

Taken together, our results identify targets for modifier screens that address the contribution of TAH-DEGs to tumor development using both classical and molecularly induced tumor mutants.

Additionally, our work points toward a potential communication network that links TAHs, SG tumors, and non-tumor tissues (**Figure 8**). This likely includes efferocytosis of apoptotic bodies derived from hyperplastic SGs (18), leading to an extended presence of SG transcripts (passenger transcripts) in TAHs before their degradation. While most of the DEGs in TAHs are genuine TAH transcripts, we find exceptions. These include two prophenoloxidases (26), which are known to be exclusively expressed in crystal cells (30) and not in the plasmatocytes we sequenced. Notably, transcripts of potential non-plasmatocyte origin also include the NF-kappaB factor Relish, a key regulator of the Imd pathway (31). While the mechanism for this cell’s non-autonomous presence of transcripts in TAHs is unknown, we hypothesize that exosomes may be likely candidates (**Figure 8, dashed arrows**). Alternatively, plasmatocytes may transdifferentiate and express transcripts that are more specific for crystal cells and lamellocytes (32,33) (**Figure 8, solid arrows).** Supporting an exosome origin, two long non-coding RNAs (CR 34335 and R 40469) we found in TAHs have been identified among the 10 most abundant non-ribosomal RNAs in exosomes released from *Drosophila* cell lines (34). Additionally, in the same study, *Arc1* was shown to be amongst the most abundant mRNAs in exosomes from one of the two cell lines used (34). Notably, in the flies’ nervous system, Arc1 is involved in the formation of capsid-like vesicles, which also contain Arc1 transcripts that are recruited through the binding of Arc1 protein to the 3’ end of the *Arc1* transcript (35). Both Arc1 and its human equivalent derive from retroviral Gag proteins (35–37) and mediate neuronal plasticity. A function for *Drosophila* Arc1 in immunity has so far not been suggested although it is expressed in non-neural tissues (FlyAtlas: (22)) and we find it is enriched in TAHs **(Figure 5)**. Of note, similar to tumor-associated macrophages, extracellular vesicles have been shown to display pro-tumor potential. Taken together, our findings on macrophage-like cells from an invertebrate provide targets that may turn out useful to steer tumor therapy even in humans (2).

## 6 Author Contribution

**UT** conceived the study. **JA** and **UT** supervised the study. **DK** experimental design, investigation, methodology and visualization **MK** performed immunohistochemistry analyses and proliferation assay as well as *Drosophila* genetics. **JA** and **DK** performed sorting and scSeq. **MM** performed mapping and computational analyses of the data. **MM**, **DK** and **MK** generated figures. **JA** supervised the computational analysis. **UT, MM** wrote the manuscript. All authors read and reviewed the manuscript.

## Acknowledgements

We thank the Microbial Single Cell Genomics and the Eukaryotic Single Cell Genomics facilities at Science for Life Laboratory (SciLifeLab), Sweden, for cell sorting as well as smart-seq2 library preparations, respectively, and the National Genomics Infrastructure, SciLifeLab for sequencing. The computations were performed using resources provided by SNIC through Uppsala Multidisciplinary Center for Advanced Computational Science (UPPMAX). We also thank Jiwon Shim (Hanyang University, Korea) for useful comments on the manuscript.

