## Supplemental data1 for "Single-cell sequencing of tumor-associated macrophages in a *Drosophila* model" (from another version of this preprint: version 1, 2023-06-18)

### 1 Supplementary Figures

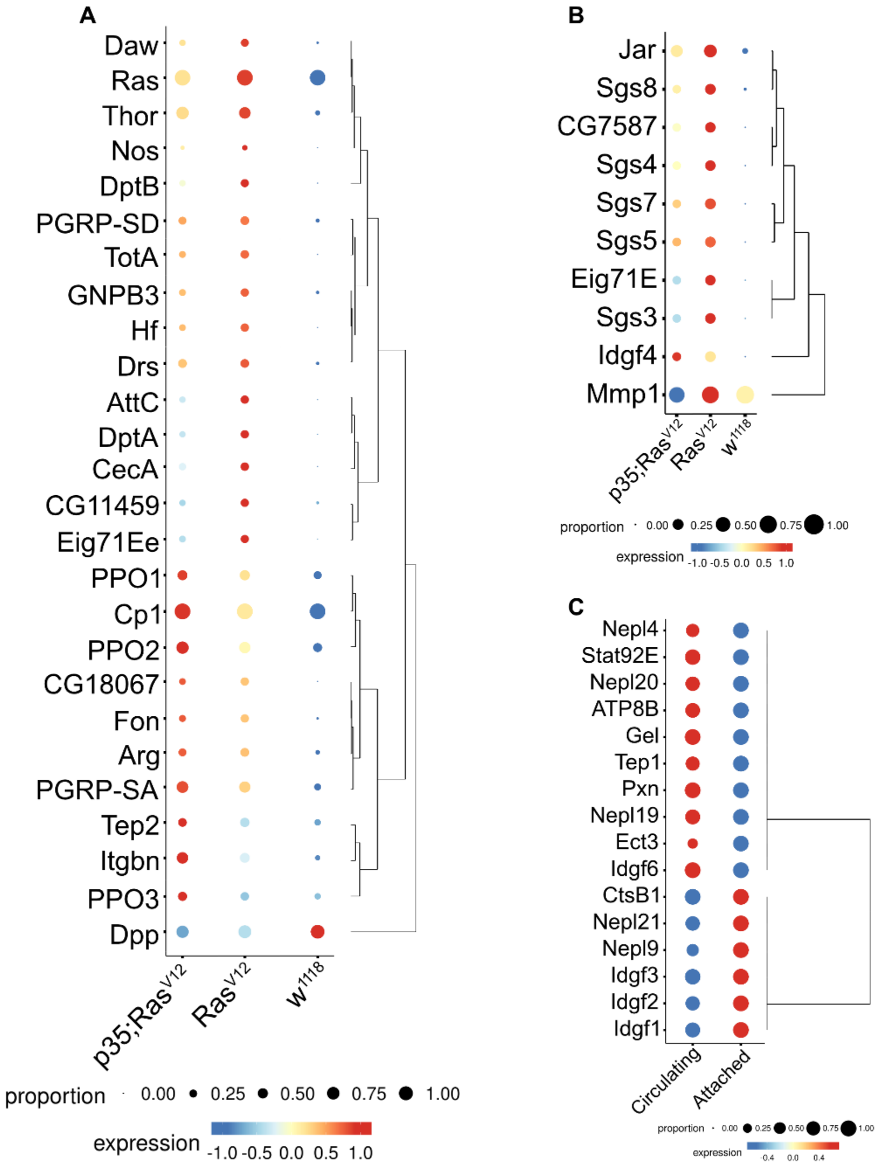

**Figure S1**

Split analysis for the genes in cluster 2 using the category Biological function which identified the terms **(A)** “Immune system process” and **(B)** “Molting process” as most significant. These two terms were compared between the three genotypes. **(C)** Genotype-specific analysis for the genes in cluster 0 in the Reactome pathway “Neutrophil degranulation” were compared between attached vs circulating cells.

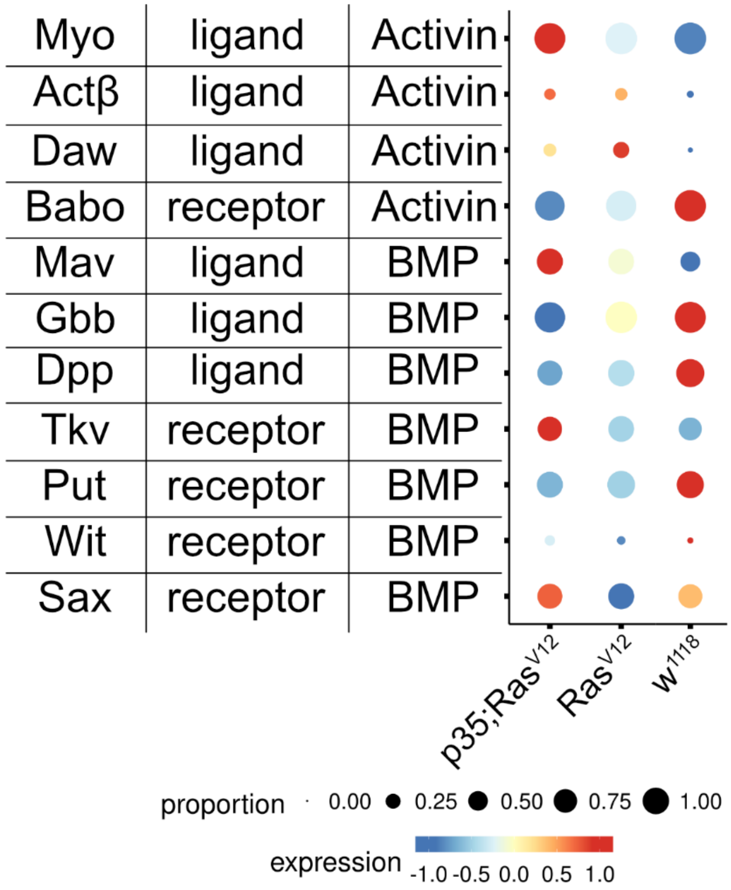


**Figure S2**

Split analysis for the genes involved in tumor necrosis factor (TGF) signaling using https://mubashe r-mohammed.shinyapps.io/Sc-drosophila/?_ga=2.6566386.119719391.1657634466-1490444630.1643316020. Both members of the activin- and BMP-like pathway are shown including ligands and receptors.
